# Classifying Irritable Bowel Syndrome Using Spatio-Temporal Graph Convolution Networks on Brain fMRI Data

**DOI:** 10.1101/2024.11.05.622062

**Authors:** Jiazhen Wu, Shuxin Zhuang, Zhemin Zhuang, Lei Xie, Mengting Liu

## Abstract

Irritable bowel syndrome (IBS) is a functional gastrointestinal disorder marked by abdominal pain and changes in stool consistency or frequency. Recent studies have explored the link between IBS and various cognitive deficits using functional MRI. Despite these efforts, an effective diagnostic or predictive model for IBS remains elusive. This shortfall is twofold: firstly, the sample sizes in these studies are typically small, and secondly, the machine learning or deep learning models currently in use fail to adequately detect the subtle and dynamic pathological changes present in fMRI data for IBS. In this study, we extracted rs-fMRI of 79 subjects with IBS and 79 healthy controls, then put them into spatio-temporal graph convolution network (ST-GCN) for classification. We also incorporated a novel interpretability module into this model to identify potential regions of interest (ROI) associated with IBS. Our model outperformed other state-of-the-art ML and DL methods with the highest average accuracy of 83.51% on our dataset. Furthermore, based on the results of our interpretability module, the Inferior Parietal Lobule (IPL.R), Inferior Frontal Orbital part (ORBinf.R), Postcentral Gyrus (PCG.R), Middle Frontal Orbital part (ORBmid.R), and Superior Medial Frontal Orbital part (ORBsupmed.L) were identified as top 5 important brain regions for distinguishing IBS patients from the control group, which are consistent with the brain regions identified in previous literature reviews. We also conducted an external data-driven experiment to further validate the effectiveness of the interpretability module. The results indicate that the selected regions significantly impact IBS.

## 1. Introduction

Irritable bowel syndrome (IBS) is a functional gastrointestinal disorder characterized by abdominal pain coupled with alterations in stool form or frequency. The prevalence of the disease ranges from 5% to 10%, primarily affecting young and middle-aged individuals. For most patients, IBS commonly manifests as a cycle of recurrence and remission (Li et al., 2021a; Ma et al., 2015; Yu et al., 2022). Owing to its persistent and intermittent nature, IBS can significantly impact patients’ quality of life and occupational productivity (Qi et al., 2016b).

Majority of studies have supported that IBS patients have high level of neuroticism and exhibit neurological deficiencies (Padhy et al., 2015). IBS is therefore considered as a new pathological approach as a brain-gut axis disorder (Balmus et al., 2020). In recent years, many studies have investigated the association between IBS and various brain impairments. Based on the findings of the survey (Farmer et al., 2012), the majority of collected articles indicate that patients with IBS often experience long-term chronic pain. This might result in the alterations in functional activity of various brain regions, along with the atrophy of gray matter and the compromised integrity of white matter in the brain.

Functional magnetic resonance imaging (fMRI) stands as a pivotal modality for unraveling the intricacies of cerebral function, comprising two primary approaches: resting-state fMRI and task-based fMRI. Task-based fMRI examines the brain’s activity in response to specific stimuli or cognitive challenges; however, it often incorporates confounding variables such as fluctuations in participant motivation and variations in task engagement. In contrast, resting-state fMRI presents a distinctive advantage by delineating the brain’s inherent connectivity patterns within a non-demanding context. For the above reasons, resting-state functional magnetic resonance imaging has been applied in the investigation of a wide range of brain disorders such as autism, Alzheimer’s disease and alcoholism. Due to its outstanding spatio-temporal properties, resting-state fMRI has demonstrated remarkable utility across diverse research domains, including neural function localization, brain network delineation, and exploration of implicated brain regions in various neurological disorders. In recent years, many researchers have started utilizing resting-state fMRI to explore the brain function of IBS. A review (Yu et al., 2022) in 2022 summarized 22 studies conducted on IBS disease using fMRI data analysis. This review highlighted alterations in brain networks in fMRI among IBS patients, predominantly concentrating in regions implicated in visceral sensation, emotional processing and pain modulation, ascertained through a comprehensive summary of previous literature. These studies show that the chronic effects of IBS lead to changes in brain function in patients that differ from those observed in healthy individuals. This serves as a pivotal basis for the application of machine learning and deep learning in the classification of IBS using fMRI data from the brain.

In recent years, a surge in the application of supervised machine learning approaches has been observed in the classification and analysis of IBS. In this study (Nan et al., 2020), researchers investigated changes in the postcentral cortex of patients with irritable bowel syndrome by extracting the regional homogeneity (ReHo) of the postcentral cortex (poCC) and the average ReHo of all voxels in the whole brain, along with the seed-based correlation based on the four seed regions: left poCC, right poCC, left insula and right insula, as inputs to a Support Vector Machine (SVM) classifier. The classification results showed excellent performance of the ReHo of poCC and seed-based correlation map (AUC>0.7). In this study (Mao et al., 2020), the researchers classified IBS and control groups using the resting-state functional connectivity (rs-FC) of the habenula-dlPFC and habenula-thalamus as input features for SVM classifier, achieving a classification accuracy of 71.5%. In this study (Lillebostad, 2019), the researchers extracted functional connectivity matrices from rs-fMRI using Pearson’s correlation and partial correlation, and utilized them as input for six machine learning models for classification, ultimately achieving an overall accuracy of 55%. In general, the application of machine learning models for the classification of IBS does yield some effectiveness, but it also encounters many challenges. For instance, in machine learning models, researchers often rely on predefined specific brain regions or functional connectivity between them as features for the classification model. However, as the brain is an integrated whole, its various parts exhibit complex interactions, thus relying solely on the variations in a few brain regions as criteria for judgment is evidently insufficient. Furthermore, previous studies have been limited by small sample sizes, with the largest sample database involving only 68 participants, including both IBS and control. Additionally, it is apparent that the accuracy results obtained from machine learning are relatively low, failing to accurately differentiate between IBS patients and control groups.

Deep learning’s principal divergence from traditional machine learning lies in the process of feature extraction. While conventional machine learning necessitates the manual extraction of features considered to significantly impact classification results, deep learning follows an autonomous feature extraction approach. In recent years, with more abundant computational resources available, deep learning models that gradually abstract features from input data through multi-layer neural network structures have been increasingly applied to the classification of various mental disorders, such as autism spectrum disorder (ASD) (Bedel et al., 2023; Dvornek et al., 2017; Hao et al., 2024; Heinsfeld et al., 2018; Li et al., 2021b; Wang et al., 2021), mild cognitive impairment (MCI)(Meszlényi et al., 2017; Suk et al., 2016), and attention deficit hyperactivity disorder (ADHD)(Sims, 2022; Zhang et al., 2022), and many other brain diseases(Huang et al., 2023; Liu et al., 2022; Tang et al., 2023; Zeng et al., 2024; Zhang et al., 2022). Some studies have used deep learning models for classification of IBS (Hachuel et al., 2019; Tabata et al., 2023; Wiriyasuttiwong et al., 2023). In this study (Hachuel et al., 2019), researchers collected 938 images of human feces from anonymous sources to investigate changes in fecal morphology caused by functional gastrointestinal disorders, including irritable bowel syndrome, chronic constipation, and chronic diarrhea. They used ResNet to classify these images into three categories: “constipation”, “normal” and “diarrhea”, achieving an average accuracy of 74.26% on a test set containing 272 images. In this study (Tabata et al., 2023), researchers collected colonoscopy images from IBS patients (n=35) and asymptomatic healthy subjects (n=88) and employed Google Cloud Platform AutoML Vision, resulting in a positive predictive value, precision, and recall of the model being 81.2%, 72.6%, and 72.6%, respectively. Unfortunately, to the best of our knowledge, there is currently no research on classifying IBS using resting-state fMRI data, which is also the driving force behind our study.

Given that fMRI data contains rich temporal and spatial information, it’s noteworthy that functional connectivity exist not only spatially between brain regions but also manifest through temporal fluctuations. To comprehensively characterize the spatio-temporal information of the brain, we have chosen to use the spatio-temporal graph convolutional network (ST-GCN) as the classification model. Compared to other deep learning models that primarily focus on static analysis (i.e., using static data such as functional connectivity matrices or graph theory attributes as inputs), ST-GCN can extract features sequentially from spatial and temporal dimensions, allowing it to focus on the dynamic development of the disease throughout the entire fMRI process. It better leverages the advantage of fMRI data that having a temporal dimension compared to other data format.

Although the earlier paragraph outlined several advantages of deep learning compared to traditional machine learning, due to their non-linear underlying structures, most deep learning models are commonly labeled as “black boxes”, and the interpretability and reliability of models’ predictions remain unclear (Castelvecchi, 2016). In the clinical settings, the absence of interpretability in decision-making processes is subject to critique (Marcus, 2018; Shortliffe and Sepúlveda, 2018), leading to a relatively low practical acceptance of deep learning methods in the medical field. Recently, various interpretable methods have been introduced for deep learning, such as post-hoc saliency-based methods like SHAP (Lundberg and Lee, 2017), gradient-weighted methods like grad-CAM (Selvaraju et al., 2017), and layer-wise backpropagation techniques (Bach et al., 2015), etc. For example, in previous experiments involving the classification of fMRI data, BrainGNN (Li et al., 2021b) added an interpretability module to its model, identifying the important brain regions that the model believed had the greatest impact on classification results. However, the interpretability module of BrainGNN is not suitable for ST-GCN because it fails to account for the dynamic changes in functional connectivity over time. In this study, in order to identify brain regions that have undergone functional changes due to long-term effects of IBS, we independently designed a new interpretability module based on the construction of the ST-GCN model to adapt to our own classification experiments. Ultimately, we identified brain regions highly influenced by the IBS disease.

In summary, to address the aforementioned requirements and concerns, this study employs a ST-GCN to classify IBS using a dataset of 79 IBS patients and 79 control subjects. It should be clarified that the fMRI data included in this dataset is resting-state fMRI and is cross-sectional in nature. This represents a significant increase in sample size compared to previous studies. Additionally, we have integrated an interpretability module specifically designed for ST-GCN. This module aims to fully utilize the temporal and spatial data in the rs-fMRI, with the goal of identifying brain regions that are clinically significant in detecting IBS impairments in the human brain.

## 2. Methods and Materials

### 2.1. Participants

This study included a total of 158 subjects (79 patients with IBS-D and 79 healthy controls), 79 patients diagnosed with IBS-D (abbreviate as IBS below) were enrolled in the Department of Gastroenterology of the First Affiliated Hospital of Shantou University Medical College from September 2021 to May 2023. Patients meeting the Rome IV diagnostic criteria for IBS, possessed an educational background of at least 9 years, fell within the age range of 18 to 50 years, and had not utilized antibiotics, selective serotonin reuptake inhibitors, or opioids within the preceding 3 months were included. 79 healthy controls (HC) matching the IBS group in age, sex, and education level were enrolled at the same time. Exclusion criteria: (1) previous history of organic gastrointestinal diseases such as inflammatory bowel disease, colon polyp and gastrointestinal surgery; (2) participants who are allergic or have taken probiotics, painkillers and other drugs in the past half month; (3) brain imaging examination found craniocerebral lesions or any neuropsychiatric diseases related to cognitive dysfunction and infectious or metabolic diseases (such as Alzheimer’s disease, multiple sclerosis, epilepsy, diabetes, etc.); (4) participants with serious mental spectrum diseases; (5) severe acute/chronic cardiac, liver and renal dysfunction; (6) patients with contraindications to MRI scanning.

This study was approved by the Ethics Committee of the First Affiliated Hospital of Shantou University Medical College (NO.B-2021-235). All subjects were informed of relevant research matters and signed written informed consent.

### 2.2. Image data acquisition and preprocessing

The resting state fMRI data were performed using GE 1.5T MR scanner at The First Affiliated Hospital of Shantou University Medical College. The rs-fMRI images were obtained using a T2* weighted gradient echo-echo plane imaging technique (GRE-EPI sequence), the imaging parameters were as follows: TR = 2000ms, TE = 45ms, FA = 90°, matrix size = 64×64, FOV = 250mm×250mm, and slice thickness = 6.0mm. The acquisition included 20 interleaved axial slices.

The rs-fMRI data was preprocessed using the GRETNA toolbox (http://www.nitrc.org/projects/gretna/). The preprocessing steps including: (1) eliminate the first 10 time point images; (2) slice timing correction; (3) head movement correction: data of subjects whose translation≥2mm and rotation≥2° were excluded; (4) spatial standardization: the fMRI images of all subjects were normalized to the standard template of the Montreal Neurological Institute (MNI152) space; (5) regression covariates: regression interference parameters to remove the influence of covariates such as noise and cerebrospinal fluid; (6) spatial smoothing: the space smoothing process is carried out by gaussian kernel with half height and full width of 6mm. (7) the regression of nuisance covariates was performed to remove noisy signals from the CSF and WM.

After preprocessing the image data, we used Automated Analytical Labeling (AAL) to determine 90 ROIs and extracted the average BOLD signal time series of all ROIs, and conduct Pearson correlation analysis one by one. In order to make the FC values follow a more normal distribution, we further processed the data using the fisher-z transform.

### 2.3 Network architecture of ST-GCN

The overall network structure is clearly illustrated in Figure.1, where three ST-GC blocks with different channel sizes were concatenated to enhance the extraction of spatial-temporal characteristics. The features convolved through three ST-GC blocks will be fed into average pooling and fully connected layers to obtain the final classification results.

#### 2.3.1 Definition of Spatio-temporal Graph

We consider the time series of each participant as a spatio-temporal graph *G* = (*V, E*). From a spatial perspective, it is regarded as a spatial graph *G*_*s*_ at each time point, the nodes of which are fully connected, each node represents a brain region, with the node features being the blood oxygen concentration signals of the corresponding brain regions at that time. The edge weights of the spatial graph *G*_*s*_ are consistently constructed across all participants, the time series data of all the participants are concatenated along the time dimension, resulting in a long time series for each brain region. Then this long time series was utilized to calculate the Pearson’s correlation between brain regions according to brain atlas AAL90. In other words, the edge weight matrix obtained through this method is not influenced by time. Therefore, during our temporal convolution operations, the graph’s edges are not taken into account; instead, we rely solely on the features of the nodes over time. This adjacency matrix is then utilized in multi-scale spatial graph convolution operators to incorporate spatial information into the graph.

#### 2.3.2 Spatial-temporal graph convolution (ST-GC) block

Based on the (Gadgil et al., 2020), the formulation of a ST-GC block is as follows:

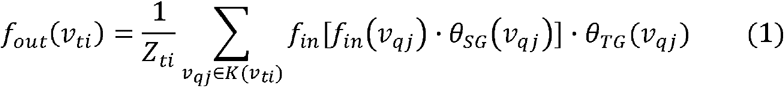

Where *v*_*ti*_ = (*t* = 1,2, …, *T*; *i* = 1,2, … *N*) represents a node, *t* denotes the time point, and *T* represents the width of the time window. Subscript *i* represents the number of brain regions, and *N* represents the total number of brain regions in the brain atlas. *Z*_*ti*_ represents a normalization factor. *K* (*v*_*ti*_) denotes the set of neighboring nodes to *v*_*ti*_, encompassing all adjacent nodes both in space and time domain. *v*_*qj*_ ∈ *K* (*v*_*ti*_) denotes the neighboring nodes of 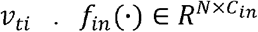 represents the input node features, where *C*_*in*_ denotes the number of input node feature channels, and 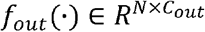 represents the output node features, *C*_*out*_ denotes the number of output node feature channels. Through the formulation (1), the features of the nodes within *K* (*v*_*ti*_) are sequentially convolved by spatial convolutional kernel 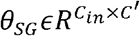, and temporal convolution kernel 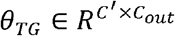 in turn, finally aggregating the features of the neighboring nodes in the set *K* (*v*_*ti*_).

Fig. 1(a) illustrates the internal structure of a ST-GC Block, where the input spatio-temporal graph data passes through two ways: one performing sequential multi-scale spatial graph convolutions and multi-scale temporal convolutions, and the other conducting spatial-temporal fusion convolutions. Subsequently, after feature fusion and a multi-scale temporal convolution module, the data flows into the next ST-GC block. Fig. 1(b) demonstrates the structure of multi-scale spatial graph convolution, where we compute eight scales of adjacency matrices based on an original adjacency matrix *A*, representing the relationships between different brain regions at various distances. After obtaining these adjacency matrices, we perform multi-scale graph convolution on the input spatial graph and output the results. Fig. 1(c) illustrates the process of spatial-temporal fusion convolutions. The structure of this block is similar to multi-scale spatial graph convolution, after multi-scale graph convolutions, the data undergoes a 3D convolution block to further fuse spatio-temporal information. Fig. 1(d) showcases the structure of multi-scale temporal convolutions, where features at different time scales are integrated by setting different dilation values.

**Fig. 1.**
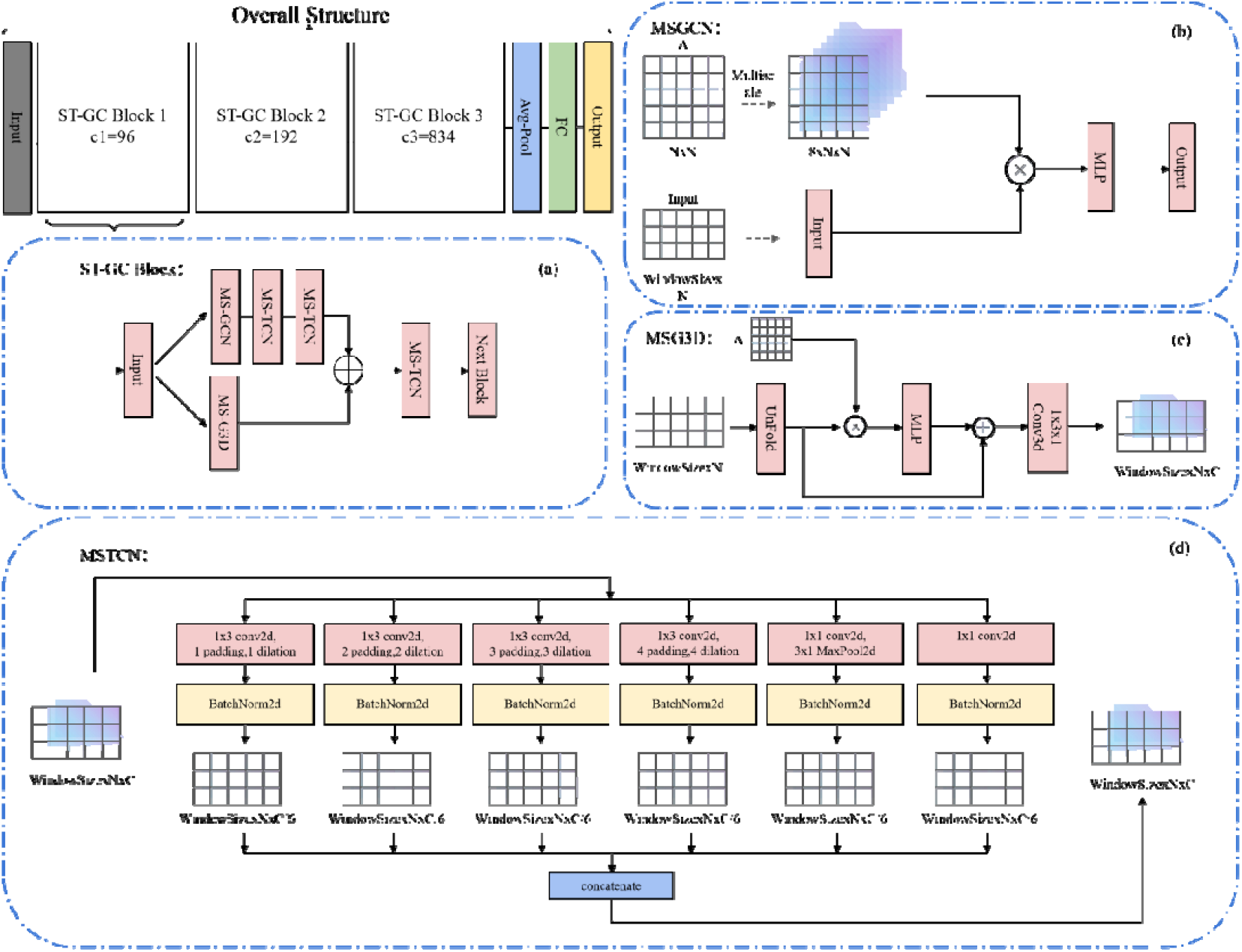
Overall Structure of the ST-GCN: Our overall network architecture consists of three ST-GC blocks with different channel numbers and a final fully connected layer. Figure (a) illustrates the internal structure of the ST-GC block, while figures (b), (c), and (d) respectively depict the detailed network architectures of MSGCN, MSG3D, and MSTCN within the ST-GC block.

#### 2.3.3 Multi-scale Spatial Graph Convolution Network (MSGCN)

The MSGCN consists of modules for graph convolution with input data and adjacency matrices at multiple scales, as well as multilayer perceptron. The classic spectral graph convolution method was applied for multi-scale convolution of the spatial graph, referred as, using a specific convolution kernel. To minimize computational complexity, we adopted the strategy outlined in the GCN paper (Karypis and Kumar, 1998), which involves using the Chebyshev Polynomials Approximation. This approach limits the convolution kernel to polynomials of the eigenvalues, denoted by, of the Laplacian matrix corresponding to the edge weight matrix. This Laplacian matrix has been normalized through the Symmetrically Normalized Graph Laplacian technique to produce a symmetric matrix. The filtering operation is represented as the multiplication of the kernel and the signal transformed by the graph Fourier transform, followed by altering the output channel number through a multi-layer perceptron. Meanwhile, by adjusting the number of terms in the polynomial, i.e., the highest degree of in the polynomial, we modify the spatial graph convolution’s receptive field (in this experiment, 8 different spatial scales are utilized) to enable more extensive range and deeper level information propagation and feature aggregation, thereby enhancing the feature extraction and representation capability in the GCN. The formulation of the spatial graph convolution transformed into spectral graph convolution is as follows:

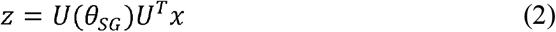

Where 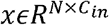 represents the input features and its dimensions, 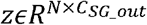 represents the output features and its dimensions, *U* ∈ *R*^*N*×*N*^ denotes the matrix of eigenvectors of the normalized graph Laplacian 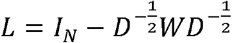, *I*_*N*_ is the identity matrix with dimensions consistent with the number of brain regions, and is the degree matrix. Through the spectral graph convolution, we have altered the channel numbers of the features of nodes within different neighborhood ranges in the input spatial graph, enabling a richer integration of information from adjacent nodes within the spatial domain.

#### 2.3.4 Multi-scale Temporal Convolution Network (MSTCN)

MSTCN is composed of multi-scale time-dimensional convolution modules and common 2D convolution modules. Following the adjustment of the node feature channel numbers through multi-scale spatial convolution, we perform temporal convolution on the convoluted graph at multiple scales. We utilize one-dimensional convolutional kernels to convolve the feature vectors from every single brain region along the temporal dimension. At each scale, we modify the dilation rates or the size of the temporal kernel to progressively widen the field of view of the temporal domain convolutional kernel along the temporal dimension. Additionally, we incorporate residual connections to address the gradient vanishing problem and improve training efficiency. Finally, by concatenating the results of multiple temporal convolutions at different scales, we obtain a representation of fMRI that integrates multi-scale spatial features and multi-scale temporal features.

#### 2.3.5 Multi-scale Spatio-temporal Convolution Network (MSG3D)

MSG3D is a module for spatio-temporal feature fusion. In addition to sequential convolutions in both spatial and temporal domains on fMRI data, the model also incorporates spatio-temporal fusion convolutions on fMRI data. This block replicates and concatenates the original adjacency matrix *A* to generate a 3*N* × 3*N* adjacency matrix, and the undergoes the same multi-scale operations as in MSGCN. This process yields multi-scale adjacency matrices shared by adjacent three time points, enabling the aggregation of spatial graphs from the three time points for graph convolutions. Subsequently, a 3D convolution is applied to the feature matrix after multi-scale spatio-temporal graph convolutions to further integrate the spatio-temporal information from these three time points.

#### 2.3.6 Training Details and Hyperparameters

In this project, we conducted a five-fold-cross-validation on rs-fMRI from 79 IBS patients and 79 control subjects to classify the disease status. In our five-fold-cross-validation experiment, we allocated 60% of the data as training data, 20% as validation data, and the remaining 20% as testing data. By inspiration of (Gadgil et al., 2020), we experimented with different window sizes, ranging from 30 to 160, to explore the optimal classification results. We empirically set the spatial scale *K*_*s*_ to 8; for the multi-scale temporal graph convolution module, the time scale *K*_*T*_ was set to 4, as this optimally combines higher-order spatial features with temporally distant features. Additionally, in this experiment, we only used an NVIDIA RTX A6000 graphics card to complete all related experiments.

### 2.4 Comparative experiments

#### Compare with deep learning models

In the comparative experiment, we applied representative deep learning models including graph convolutional network (GCN), graph attention network (GAT), graph isomorphism network (GIN), BrainGNN, BolT, TapNet and TodyNet, all of which have demonstrated superior performance in public datasets. GCN is a conventional deep learning model for graph-structured data, operates by performing convolutions on graph data and utilizes the adjacency matrices and feature tensors for information propagation and feature extraction. By updating the representation of each node through the merging of neighboring node’s feature, GCN can capture the local and global topology information within the graph. GAT incorporates attention mechanisms into graph convolutional network, enabling the automatic learning of relationships between input node features, as opposed to independently updating the features of each node as done in traditional graph convolutions. GIN is a graph convolution model designed based on the injective function property of graph neural networks, with its essence lying in the belief that a powerful GNN can map isomorphic graphs to the same representation. BrainGNN, a novel application of graph convolutional networks, employs novel ROI-aware convolutional (Ra-Conv) and ROI-aware pooling (R-pool) layers, specifically designed for rs-fMRI data. BolT partitions time-series data into multiple windows and captures local features by computing attention between selected window and adjacent windows, and progressively extends local features to global features as the overlapping range of windows increases, ultimately applying them to classification. TodyNet(Liu et al., 2024) explores the dynamic spatiotemporal relationships in multivariate time series by constructing dynamic graphs. This model obtains multiple graph structures by slicing the time series into shorter segments and initializes the vertex and edge information within each segment. These segments are then input into the spatiotemporal graph model, which is based on graph isomorphism networks, for iterative processing. TapNet(Zhang et al., 2020) first embeds the multivariate time series into a one-dimensional space using LSTM and one-dimensional convolution. This embedded representation is then input into a transformer to learn prototype vectors for each class. Finally, classification is performed by comparing the embedded one-dimensional space vectors with the class prototype vectors. We employed the aforementioned deep learning models to classify our IBS dataset and compared the best classification results with ST-GCN.

#### Compare with machine learning models

Apart from the deep learning models, several machine learning models were also employed for comparative experimentation. To reduce the features and enhance the predictive power in the comparative experiment, we first employed dimensionality reduction methods on the data. These methods included principal component analysis (PCA), locally linear embedding (LLE), factor analysis (FA) and multidimensional scaling (MDS). For each dimensionality reduction method, different parameters were set, such as different numbers of principal components or latent factors, to achieve varying degrees of dimensionality reduction. For the PCA method, three pipelines were created: pca, pca2 and pca3, which were set to retain all principal components, half of the principal components, and one-third of the principal components, respectively. The treatment for LLE was similar to that of PCA. In the context of using the FA method for dimensionality reduction, procedures akin to those used in PCA were executed. Moreover, this approach was augmented by incorporating pipelines where the number of components was specifically set to 1000 and 2000. The approach for the MDS was the same as that for FA. Consequently, we established 16 different dimensionality reduction pipelines.

In this comparative experimentation, we employed eight different machine learning methods: linear support vector machine, gaussian naive bayes, random forest, logistic regression, lasso linear regression, k-nearest neighbors, ridge regression and multi-layer perceptron. For each of the method, exploration was conducted to discover the best classification performance by adjusting different parameters. For instance, in linear support vector machine, regularization was performed using both l1 and l2 penalty, and in the multi-layer perceptron, different hidden layers and solvers for weight optimization were set. Finally, the dimensionality reduction methods and machine learning methods were combined in various permutations, resulting in a total of 208 different machine learning classifiers. These classifiers were applied to our IBS dataset using a five-fold-cross-validation method to obtain the final classification results.

### 2.5 Interpretability Modul

We extract the weights corresponding to the best classification results from the batch during the testing process, as well as a feature tensor *X*_*best*_ that integrates spatial and temporal domains prior to entering the final fully connected layer for classification. The weights and the feature tensor *X*_*best*_ are input into the interpretability module for analysis, the process of which is detailed in Figure 2. Subsequently, we will provide a detailed description of this process.

**Fig. 2.**
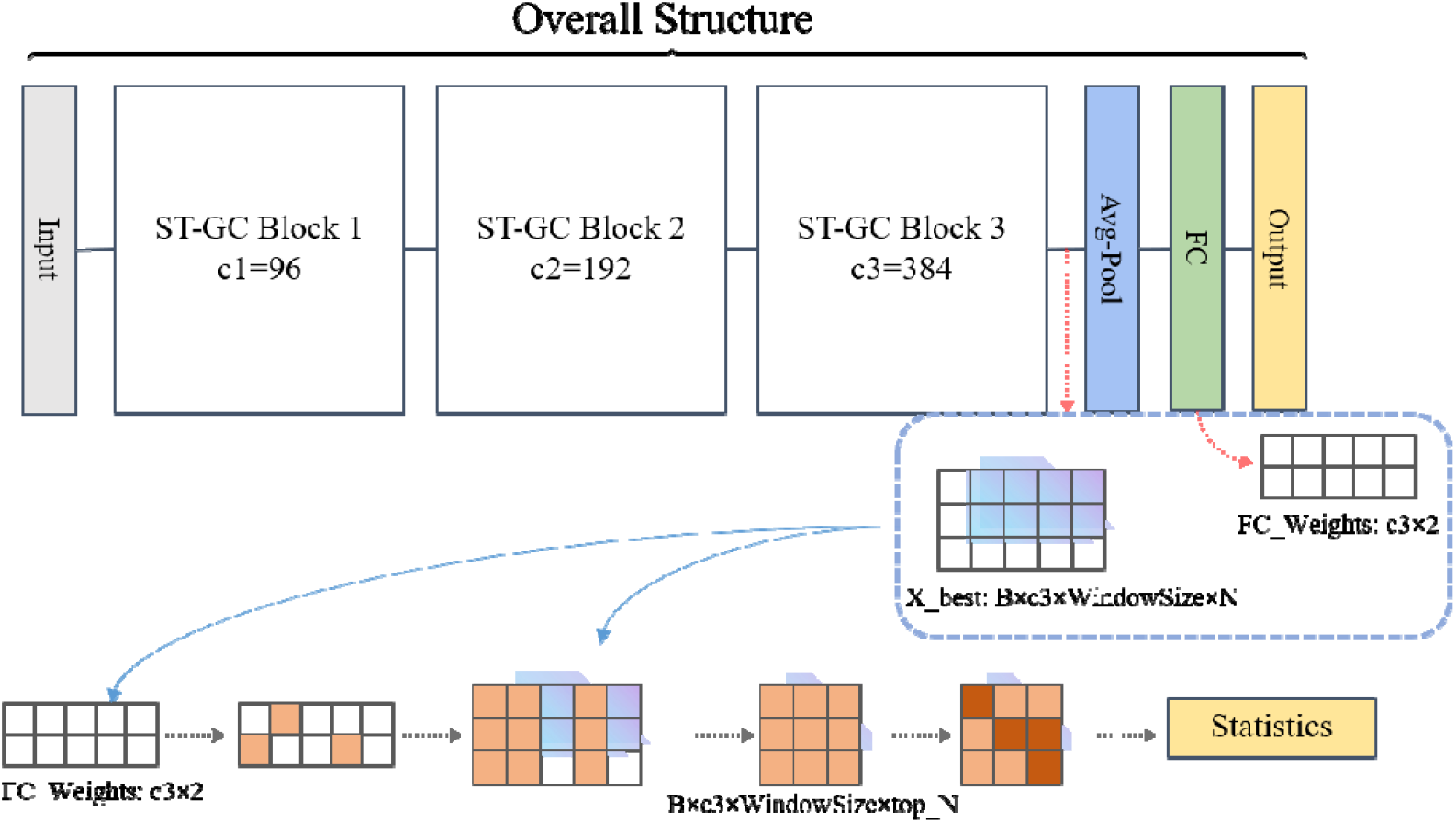
The necessary inputs for the interpretability module and the explanation process.

Initially, we extract the weights from the last fully connected layer of the neural network, denoted as FC_Weights, identifying the weights with the highest absolute values and marking their positions. We posit that these marked weights contribute most significantly to the classification outcome. Subsequently, we utilize the indices of the top-ranked weights from the previous step to extract what we consider the important channels from the feature tensor *X*_*best*_. This results in a feature vector with dimensions (B, c3, WindowSize, top_N), containing only the important channels. Next, we select the brain regions of interest from these important channels. Given that prior to entering the fully connected layer, the model flattens and averages the last two dimensions of the convolved feature tensor, we seek the elements within the dimensions (WindowSize, N) that contribute most significantly to the average value, marking their respective positions. We then count the number of marked elements in each column of the last two dimensions as a measure of the importance of that brain region, which we refer to as the importance score for each brain region. Finally, we compile and aggregate the importance scores for each brain region across all subjects in that batch, resulting in a ranked list of important brain regions.

## 3. Experiments and Results

### 3.1 Classification experiment and results on ST-GCN

Figure.3 illustrates the classification accuracy and corresponding standard deviations under varying window sizes (30-160). It is observed that the selected window size of 140 yields the optimal classification performance, achieving an accuracy of 83.51% in five-fold-cross-validation. The average accuracy across all window sizes stands at 79.03%, which significantly surpasses previous studies. However, the experimental performance was less satisfactory when the time window was either too large or too small.

**Fig. 3.**
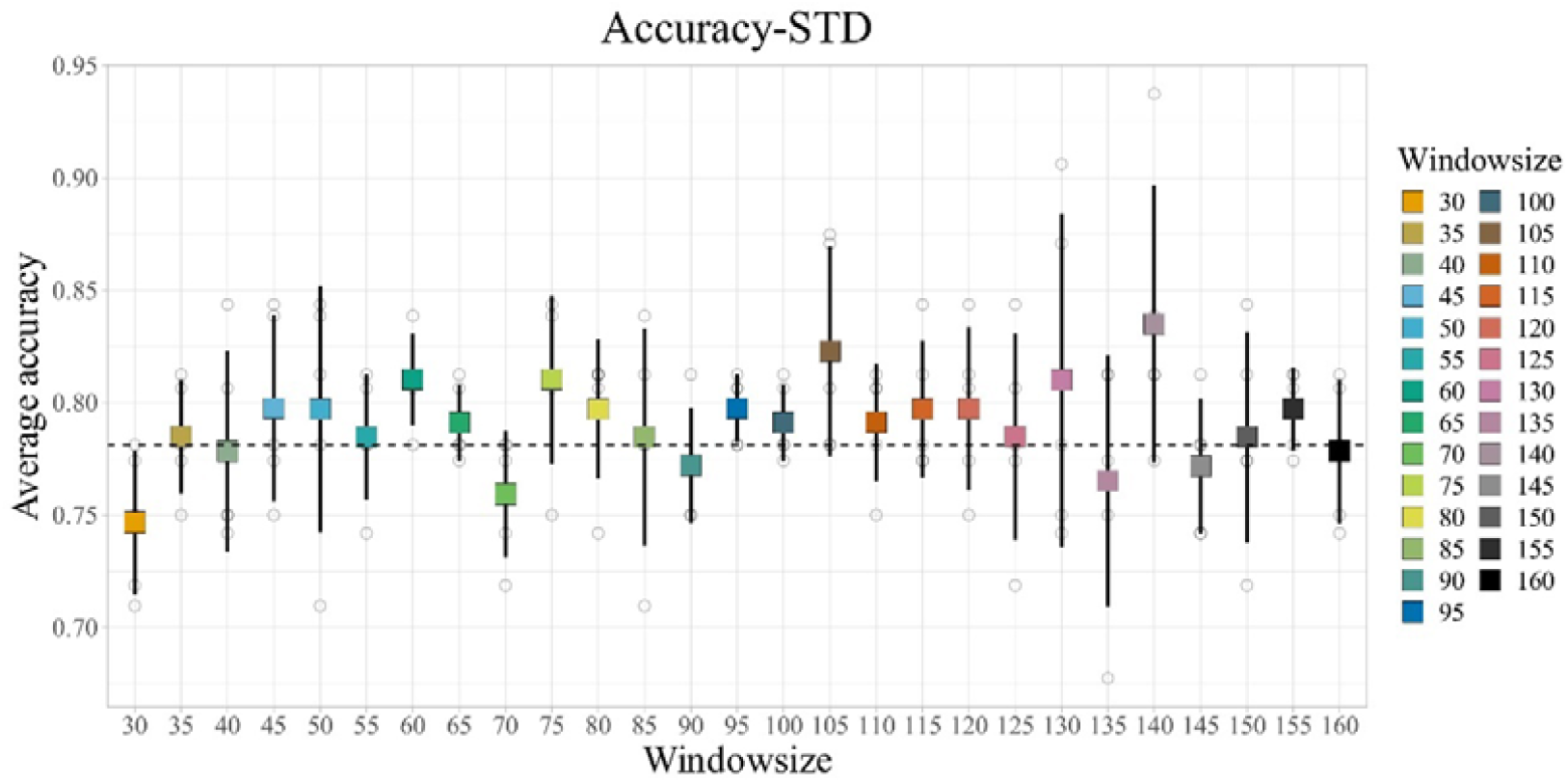
Scatter with standard deviation of average classification accuracy under different window sizes. When the window size reached 140, we achieved the optimal classification accuracy of 83.51%.

### 3.2 Comparative results

#### 3.2.1 Results with Deep Learning Models

We conducted five-fold-cross-validation experiments on seven additional deep learning models (namely GCN, GAT, GIN, BrainGNN, BolT, TodyNet, TapNet), aside from ST-GCN, while striving to optimize their hyperparameters for optimal classification performance. Table 2 displays the best classification accuracy, along with corresponding sensitivity, specificity and precision values, for these eight models. Comparative findings reveal that the classification performance of ST-GCN far exceeds that of other deep learning models, and it exhibits superior stability. We also plotted the ROC curves (Figure.4) for the best training results of these eight deep learning models to demonstrate the stability of ST-GCN.

**Table 1.**
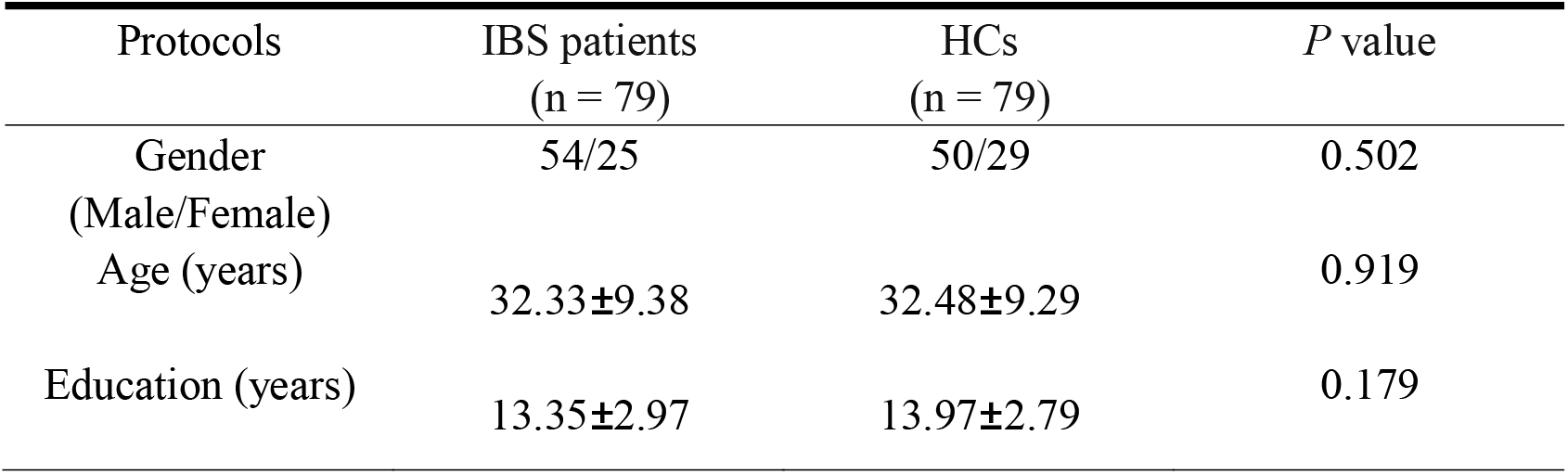
Demographics information of the IBS dataset used in this study.

**Table 2.** classification performance of deep learning models.

| Model | Acc mean(%)<br>( STD ) | Sens mean(%)<br>( STD ) | Spec mean(%)<br>( STD ) | Prec mean(%)<br>( STD ) |
| --- | --- | --- | --- | --- |
| GCN | 61.98(0.014) | 57.42(0.010) | 66.82(0.020) | 62.78(0.017) |
| GIN | 61.74(0.018) | 52.58(0.035) | 69.58(0.022) | 64.82(0.032) |
| GAT | 58.70(0.009) | 56.18(0.015) | 61.38(0.022) | 60.06(0.010) |
| BrainGNN | 61.25(0.032) | 47.69(0.191) | 53.33(0.163) | 79.26(0.088) |
| BolT | 71.28(0.106) | 67.61(0.197) | 75.18(0.213) | 77.16(0.152) |
| TodyNet | 55.63(0.013) | 86.67(0.125) | 23.94(0.097) | 45.60(0.053) |
| TapNet | 65.00(0.046) | 90.00(0.109) | 40.00(0.188) | 61.24(0.065) |
| **ST-GCN(ours)** | 83.51(0.053) | 84.83(0.029) | 81.00(0.112) | 84.16(0.026) |

**Fig. 4.**
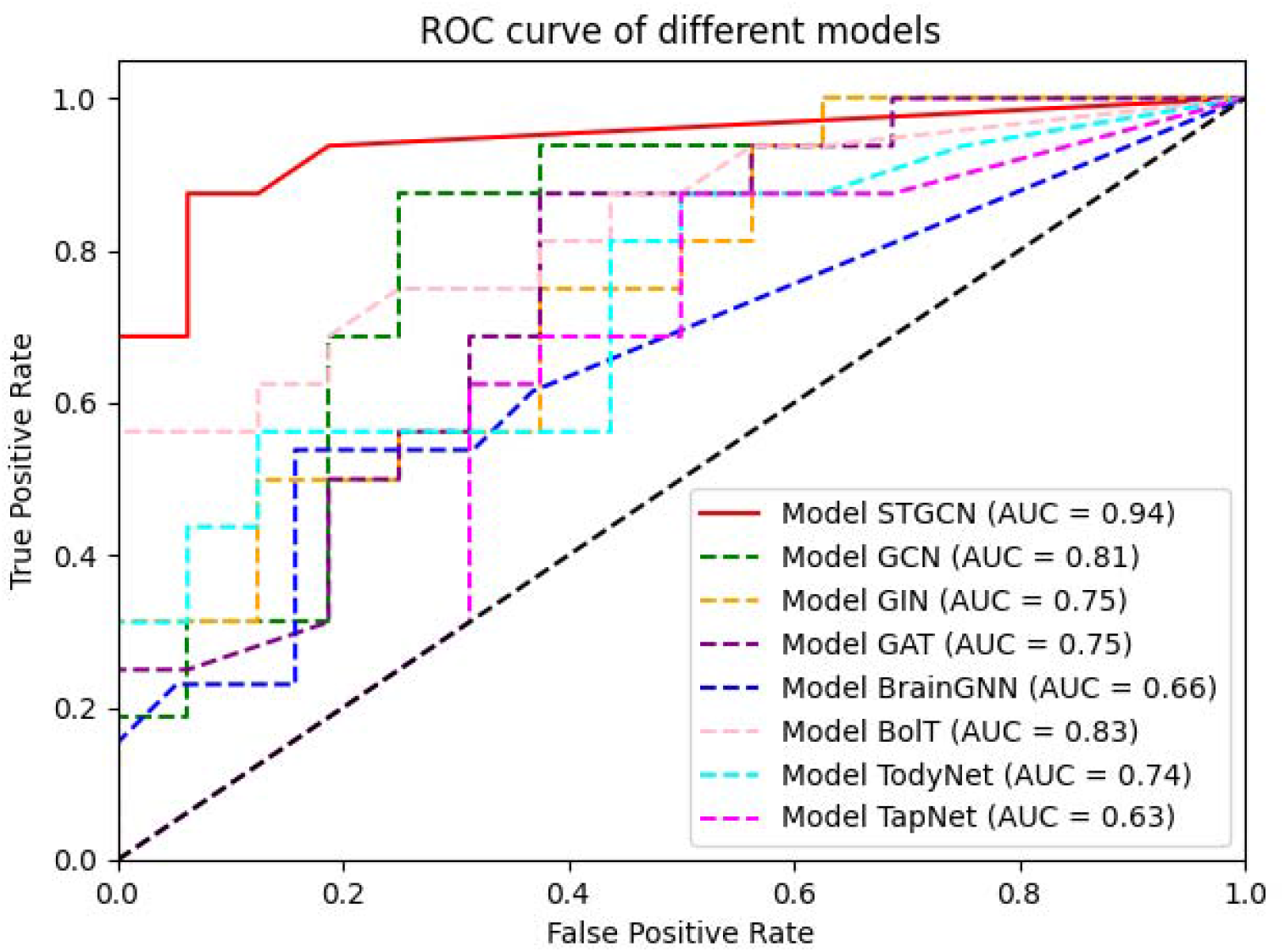
ROC curves depicting the optimal training results of eight deep learning models. For the eight deep learning models, we plotted the optimal classification accuracy for each and found that the ST-GCN model exhibited the best classification performance.

#### 3.2.2 Results with Machine Learning Models

In the comparative experiments with machine learning models, we employed 13 different methods with the best dimensionality reduction method respectively, and show the results is present in Table 3. The first column denotes the abbreviation of the utilized pipeline, while the second column represents the average accuracy of all reduction methods for each pipeline. The third column displays the accuracy of the optimal reduction method for each pipeline. Additionally, the fourth column indicates the dimensionality reduction method employed when the best accuracy was achieved.

**Table 3.** The results of machine learning comparative experiments.

| Classifier | Average accuracy | Best accuracy | Reduction |
| --- | --- | --- | --- |
| GaussianNB | 0.4984 | 0.5688 | MDS |
| RandomF | 0.5789 | 0.6625 | LLE |
| anova_svcl1 | 0.5297 | 0.6313 | LLE |
| anova_svcl2 | 0.5348 | 0.6438 | LLE |
| knn | 0.5699 | 0.6500 | FA2/PCA/PCA2 |
| logistic_l2 | 0.5645 | 0.6438 | LLE2 |
| neural10 | 0.5809 | 0.7000 | LLE2 |
| neural10a | 0.5855 | 0.6438 | LLE |
| neural5 | 0.5695 | 0.7000 | LLE2 |
| neural5a | 0.5148 | 0.6250 | FA2 |
| ridge | 0.5621 | 0.6438 | LLE2 |
| svc_l1 | 0.5590 | 0.6500 | LLE2 |
| svc_l2 | 0.5711 | 0.6438 | LLE2 |

The classifiers in the first column are: (1)gaussian naive bayes; (2)random forest; (3)linear support vector machine with l1 penalty and ANOVA feature selection method; (4)linear support vector machine with l2 penalty and ANOVA feature selection method; (5)k-nearest neighbors; (6)logistic regression with l2 penalty; (7)10 hidden layers multi-layer perceptron; (8)10 hidden layers multi-layer perceptron with Adam solver; (9)5 hidden layers multi-layer perceptron; (10)10 hidden layers multi-layer perceptron with Adam solver; (11)ridge regression; (12)linear support vector machine with l1 penalty; (13)linear support vector machine with l2 penalty.

In our machine learning pipelines, employing a multi-layer perceptron (MLP) classifier which configuring 10 hidden layers, yielded the highest average accuracy of 58.55%. Additionally, utilizing LLE with half the size of the training dataset as the reduced feature dimension, both 10-hidden-layer and 5-hidden-layer MLP classifiers achieved the highest accuracy of 70%. Comparatively, traditional machine learning methods exhibited significantly lower average accuracy compared to the ST-GCN model.

#### 3.2.3 Results of ablation experiments

We conducted ablation experiments on all submodules within the ST-GC blocks simultaneously to verify the necessity of each submodule. Specifically, we performed ablation on MS-G3D, MS-GCN, and three MS-TCNs, resulting in a total of five distinct ablation experiments for validation. Table 4 presents the results of the five ablation experiments conducted using five-fold-cross-validation.

**Table 4.** The results of ablation experiments.

| Ablated submodule | Acc mean(%)<br>( STD ) | Sens mean(%)<br>( STD ) | Spec mean(%)<br>( STD ) | Prec mean(%)<br>( STD ) |
| --- | --- | --- | --- | --- |
| MS-GCN | 81.71(0.072) | 83.67(0.074) | 79.83(0.127) | 81.61(0.087) |
| MS-G3D | 81.67(0.045) | 88.58(0.073) | 74.83(0.103) | 78.51(0.061) |
| MS-TCN-1 | 81.65(0.023) | 81.00(0.105) | 82.33(0.133) | 83.86(0.075) |
| MS-TCN-2 | 82.28(0.025) | 82.17(0.077) | 82.33(0.072) | 83.03(0.053) |
| MS-TCN-3 | 79.78(0.023) | 79.66(0.051) | 79.83(0.091) | 80.80(0.068) |

#### 3.2.3 Results of different hyperparameters

Firstly, we sought the optimal hyperparameters by adjusting the spatial scale parameters of the MS-GCN and MS-G3D modules (they are denoted as gcn_scales and g3d_scales in table 5 respectively), i.e., modifying the receptive field in the spatial convolution process. Subsequently, we adjusted the temporal scale parameters (which denoted as tcn_scales in Table 5) by progressively increasing the dilation range of the temporal convolutional kernel to observe the classification performance under different temporal receptive fields. Table 5 shows the classification results obtained under different hyperparameter configurations.

**Table 5.** The results of different hyperparameters.

| gcn_scales | Acc<br>mean(%)<br>( STD ) | g3d_scales | Acc<br>mean(%)<br>( STD ) | tcn_scales | Acc<br>mean(%)<br>( STD ) |
| --- | --- | --- | --- | --- | --- |
| 1 | 77.82(0.053) | 1 | 77.82(0.022) | 1 | 78.49(0.053) |
| 2 | 81.61(0.026) | 2 | 77.80(0.042) | 2 | 75.95(0.043) |
| 3 | 72.12(0.027) | 3 | 77.84(0.057) | 3 | 76.61(0.041) |
| 4 | 79.15(0.041) | 4 | 79.74(0.068) | 4 | 83.51(0.053) |
| 5 | 77.90(0.054) | 5 | 79.11(0.033) | 5 | 81.01(0.045) |
| 6 | 76.57(0.076) | 6 | 78.45(0.059) | 6 | 75.97(0.036) |
| 7 | 77.82(0.030) | 7 | 72.78(0.033) |  |  |
| 8 | 83.51(0.053) | 8 | 83.51(0.053) |  |  |

### 3.3 Interpretability results

#### 3.3.1 Interpretability experiment and results

With the five-fold-cross-validation, we selected the five folds with a window size of 140 (Optimal classification accuracy window size) and extracted their weight matrices and final feature tensors, which were then averaged separately and input into our interpretability module to statistically assess the importance of each brain region. Figure.5 shows the Top 20 brain regions identified by our interpretability module as being strongly associated with IBS, and we also show the importance scores for each brain region in the figure. Results revealed that for the Inferior Parietal Lobule (IPL.R) and the Inferior Frontal Orbital part (ORBinf.R) are significantly higher than those of other brain regions. The Postcentral Gyrus (PCG.R), Middle Frontal Orbital part (ORBmid.R), and Superior Medial Frontal Orbital part (ORBsupmed.L) are in the second tiers. Subsequently, the following regions are ranked: Postcentral Lobule (PCL.R), Superior Frontal Orbital part (ORBsup.R), Caudate nucleus (CAU.R), Rectal gyrus (REC.R), Anterior and Posterior Cingulate Gyrus (ACG.R), Postcentral Lobule (PCL.L), Superior Parietal Lobule (SMG.L), Inferior Occipital Gyrus (IOG.R), Pallidum (PAL.R), Lingual Gyrus (CUN.L), Angular Gyrus (ANG.R), Supplementary Motor Area (SMA.L), Thalamus (THA.L), Hippocampus (HIP.L), Supplementary Motor Area (SMA.R). To visually assess the significance of different brain regions, we performed visualization processing on the outcomes derived from the interpretability module. For displaying cortical regions, we utilized the SurfStat toolbox (http://www.math.mcgill.ca/keith/surfstat), and for subcortical regions, we employed the Enigma Toolbox (Larivière et al., 2021).

**Fig. 5.**
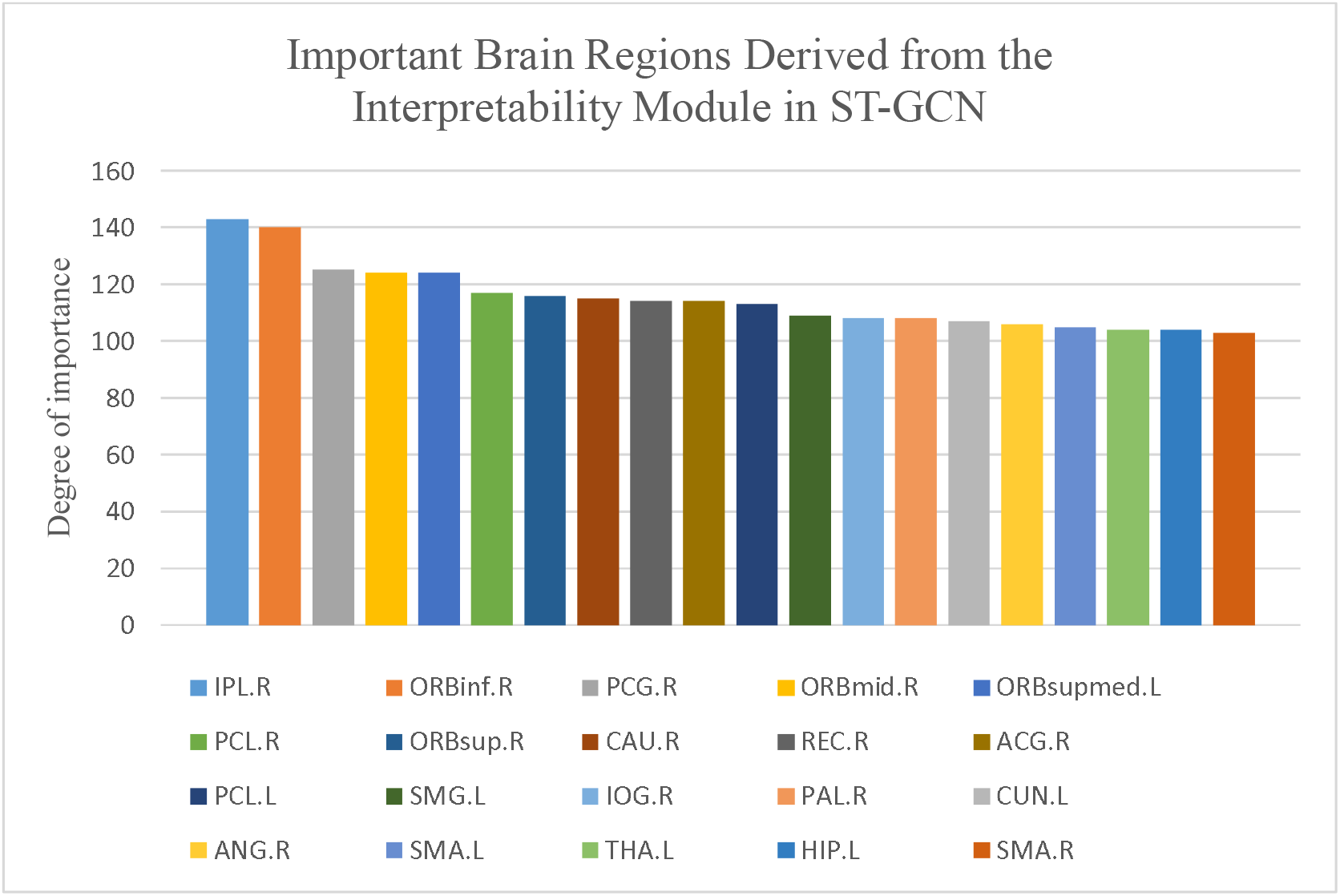
The importance scores of brain regions derived from the interpretability module. (AAL90 atlas)

**Fig. 6.**
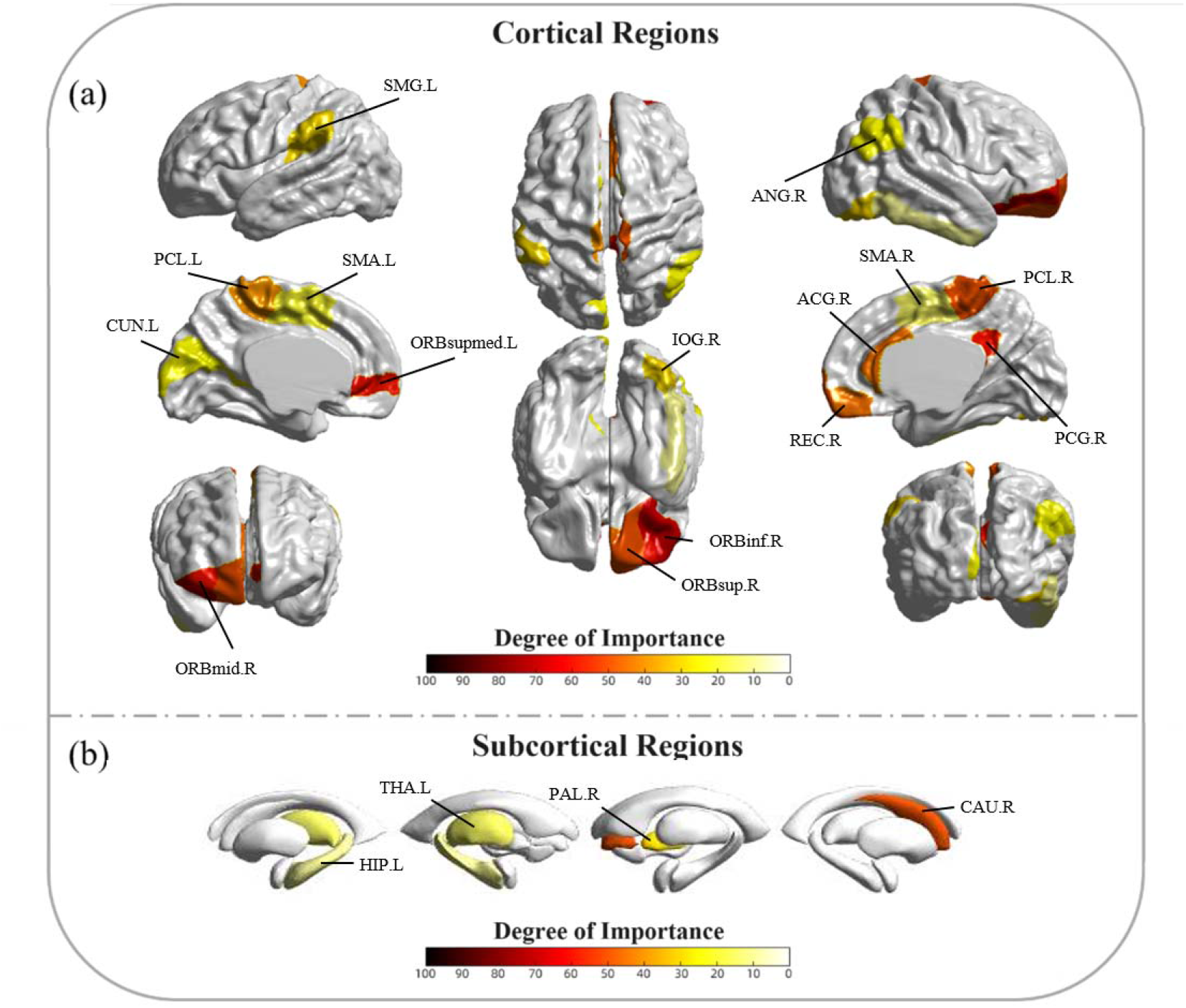
Visualization of important brain regions, with abbreviations for significant areas indicated in the figure. (a) represents crucial brain regions located on the cerebral cortex, while (b) denotes regions situated beneath the cerebral cortex. The gradient from white to red signifies increasing degree of importance.

#### 3.3.2 External validation of brain regions extracted from the interpretability module

##### Experiment only on important brain regions

To further validate the significance of our interpretability module, we extracted data from the top 20 significant regions identified by interpretability module and conducted classification experiments by inputting the time series data of these 20 significant regions into the ST-GCN model separately. Following the same experimental procedures as the whole-brain experiment, we performed five-fold-cross-validation on these data with optimal window size 140. In order to compare with the experiments described above, we also randomly selected 20 brain regions from all brain regions and conducted the same experiment (The comparative experiment will be repeated 10 times with optimal window size 140, and statistical analysis will be conducted. The brain regions selected each time will be random and mutually independent.).

The results showed that the five-fold average classification accuracy of the 20 significant regions’ data exceeded that of the 20 random regions, the top 20 significant regions exhibited an average accuracy that was close to 3% higher than the 20 random regions in window size 140 (p<0.001), and was comparable to the average classification accuracy of the whole-brain experiment in several window sizes. This demonstrates the validity of the conclusions drawn by the interpretability module. The table 6 below shows the average accuracy and other binary classification attributes of five-fold-cross-validation for classification using the top 20 significant regions’ and random 20 regions’ data in time window size 140:

**Table 6.** classification performance of top 20 significant regions and random 20 regions.

| Group | Acc mean(%)<br>( STD ) | Sens mean(%)<br>( STD ) | Spec mean(%)<br>( STD ) | Prec mean(%)<br>( STD ) |
| --- | --- | --- | --- | --- |
| Random 1 | 75.30(0.043) | 77.17(0.034) | 73.25(0.098) | 75.15(0.064) |
| Random 2 | 75.28(0.074) | 78.75(0.151) | 72.08(0.124) | 74.84(0.095) |
| Random 3 | 74.58(0.083) | 70.92(0.187) | 78.75(0.146) | 80.00(0.130) |
| Random 4 | 75.30(0.059) | 77.50(0.129) | 73.42(0.114) | 75.54(0.086) |
| Random 5 | 73.95(0.086) | 74.50(0.087) | 73.25(0.126) | 74.46(0.110) |
| Top 20 | 78.51(0.057) | 81.25(0.040) | 75.67(0.082) | 77.73(0.067) |

**Table 7.** top 10 significant regions obtained by interpretability module and PBM.

| Ranking | interpretability module | PBM |
| --- | --- | --- |
| 1 | IPL.R | OLF.R |
| 2 | ORBinf.R | PAL.R |
| 3 | PCG.R | PCL.L |
| 4 | ORBmid.R | REC.R |
| 5 | ORBsupmed.L | DCG.L |
| 6 | PCL.R | ORBinf.R |
| 7 | ORBsup.R | PHG.R |
| 8 | CAU.R | PHG.L |
| 9 | REC.R | THA.L |
| 10 | ACG.R | IPL.R |

**Table 7.** top 10 significant regions’ correlation with node attributes.

| regions | IPL.R | ORBinf.R | PCG.R | ORBmid.R | ORBsupmed.L |
| --- | --- | --- | --- | --- | --- |
| node degree | √ | √ |  | √ | √ |
| closeness centrality | √ | √ |  | √ | √ |

| clustering coefficient | ✓ | ✓ | ✓ | ✓ |  |
| --- | --- | --- | --- | --- | --- |
| regions | PCL.R | ORBsup.R | CAU.R | REC.R | ACG.R |
| node degree |  |  | ✓ |  | ✓ |
| closeness centrality |  |  | ✓ |  |  |
| clustering coefficient |  |  | ✓ |  | ✓ |

##### Comparison with the Perturbation-Based Methods (PBM)

To further validate the reasonability of our interpretability module, we referenced the PBM as a comparison. PBM are a class of widely applied analytical techniques in the fields of neuroscience and machine learning. These methods involve targeted perturbations, such as input modifications or parameter adjustments, to investigate the dynamic responses and stability of neural systems or models. By employing perturbations, we mask data from one brain region at a time, calculate the distance between the predictive output of the masked data and the unmasked data, and ultimately rank the importance of brain regions from high to low based on the distances in descending order. We conducted five-fold-cross-validation experiments using a window size of 140, which yielded the best classification results, along with other optimal hyperparameters, and applied PBM to the model at the point of maximum classification accuracy. Consequently, each five-fold-cross-validation experiment produced five rankings of important brain regions, resulting in a total of 25 rankings. Finally, we statistically analyzed these 25 rankings of important brain regions to derive the order determined by PBM.

Through comparison, we found that IPL.R, ORBinf.R and REC.R appeared in the top 10 important brain regions of both interpretability methods. Additionally, within the groups of the top 20 brain regions identified by both methods, 9 regions were consistently represented in the results of both approaches. Therefore, it is evident that various interpretative methods may highlight different brain regions, and a standardized approach for evaluating these methods is currently lacking. Nonetheless, the overlapping regions identified across different techniques may represent areas of greater significance for IBS and warrant closer attention.

##### Correlation analysis

To further validate the reasonability of our interpretability module, we conducted a general linear model with gender, age and educational years as covariates between the node attributes derived from brain graphs of the patient group and the corresponding IBS-SSS (IBS symptom severity scale) scores of the patients. Specifically, we employed the functional connectivity matrix of the patient group to generate the graph and computed the degree, closeness centrality, and clustering coefficient of the graph nodes. After extracting the attributes of the corresponding brain region nodes (the top ten brain regions), we performed a linear correlation analysis between a specific node attribute of a particular brain region and the IBS-SSS scores of the patient group.

Our findings indicate a strong correlation between these three node attributes and the IBS-SSS scores of the patient group. We will discuss the analysis results of these three node attributes in turn. First, among the top ten brain regions, the node degree of IPL.R, ORBinf.R, ORBmid.R, ORBsup.R, CAU.R and ACG.R demonstrated significant positive correlations with the IBS-SSS scores (FDR corrected p’s<0.05). Next, the closeness centrality of IPL.R, ORBinf.R, ORBmid.R, ORBsup.R and CAU.R also exhibited significant positive correlations with the IBS-SSS scores (FDR corrected p’s<0.05). Finally, the clustering coefficient of IPL.R, ORBinf.R, PCG.R, ORBmid.R, CAU.R and ACG.R displayed significant positive correlations with the IBS-SSS scores (FDR corrected p’s<0.05).

## 4. Discussion

### Overall model performance

Currently, there is a lack of research utilizing deep learning for classifying fMRI data for IBS. Primarily, this is due to the limited sample size, and employing deep learning models with large parameters poses a high risk of overfitting. Through a series of experiments, supported by a large amount of data, we have demonstrated the classification effectiveness of STGCN on our IBS dataset. The model achieved an accuracy of 83.51% at a window size of 140, surpassing the accuracy of all previous classification methods while ensuring model stability. We believe that the primary benefit of the STGCN compared to traditional machine learning techniques and other deep learning models is its remarkable ability to extract and integrate spatio-temporal features from time series data for effective representation. This highlights the critical role that advanced spatio-temporal features play in enhancing the classification accuracy of brain time series data.

Furthermore, in IBS research, classification is not the primary objective, researchers are more interested in identifying biomarkers that can differentiate IBS patients from healthy subjects. Present studies often adopt a hypothesis-driven approach, whereby problematic brain regions are hypothesized and features related to these regions, such as Reho and functional connectivity, are extracted. Subsequently, machine learning models are constructed to gauge the correlation of these extracted features with IBS through classification. Despite the capabilities of existing approaches, they fall short in fully leveraging the rich spatio-temporal information available in fMRI data. In our research, we utilize the ST-GCN to thoroughly exploit the spatio-temporal characteristics of data pertinent to IBS, without presuming pathological areas. This data-driven deep learning classification approach significantly enhances classification accuracy and, by designing a comprehensive interpretability module, enables the identification of brain regions contributing significantly to classification accuracy. Because of this interpretability module, our study offers comprehensive exploration of IBS.

fMRI data is extensively utilized in brain research, not just for its high spatial resolution but also for its ability to capture temporal variations, offering insights into the brain’s dynamic activities. However, the complex spatial and temporal information in fMRI data poses a challenge for deep learning models to fully comprehend the contributions of specific brain regions over time. In this study, after fusing spatio-temporal features of fMRI data using ST-GCN, we incorporate an interpretability module designed to reverse infer neural network parameters, tackling this complexity. The brain regions pinpointed by this module have shown highly significant relevance to IBS as confirmed by our external experimental validations and literature reviews, thereby validating the efficacy of our interpretability approach. Although we classified our IBS dataset using a uniform window size and achieved optimal classification results, an important factor has been overlooked: individual differences. The individual differences arising from factors such as gender, age, educational years, pain severity, and subtype of the IBS disease cannot be ignored, even though these subjects were collected from the same medical institution. Technically, we look forward to future research that explores specific window size and optimal hyperparameters for different subgroups of IBS patients and even individual IBS patient. Clinically, we also look forward to future studies that incorporate subtype separations on IBS based on their clinical or imaging performance, this would believed to be given more nuanced information about the neuropathological factors towards the disease.

### Clinical Significance

We believe that the deep learning model applied in this study holds significant clinical implications, as it can assist physicians in improving the diagnostic accuracy of irritable bowel syndrome (IBS), thus enhancing patient treatment outcomes. Additionally, the model can effectively analyze complex patterns in brain rs-fMRI data through its interpretability module, aiding clinicians in identifying biomarkers associated with IBS, thereby facilitating earlier and more accurate diagnoses.

To translate this model into a practical diagnostic tool, we propose several steps. First, we recommend conducting multi-center clinical data collection to comprehensively validate the model’s stability and generalizability, ensuring its adaptability to diverse patient populations. Upon successful validation, we will aim to develop a user-friendly interface that seamlessly integrates into existing clinical workflows, allowing healthcare professionals to efficiently input patient data and obtain diagnostic results. We also aim to design a website where users can easily upload data and automatically receive diagnostic results. This website will protect the privacy of users’ data and assist users in making decisions. By bridging advanced machine learning techniques with clinical practice, we aspire to enhance the efficiency of the diagnostic process and ultimately improve the quality of care for patients with IBS.

### Brain regions altered by IBS

Using the brain region rankings provided by the interpretability module, we compare these identified regions to those reported in previous literature. We found that many brain regions undergoing changes belong to the default mode network, for example, the Postcentral Gyrus (PCG.R), the Anterior and Posterior Cingulate Gyrus (ACG.R) and the hippocampus (HIP.L), suggesting that the persistent pain caused by IBS is reflected in DMN. This finding is well supported by previous studies, for example, in (Farmer et al., 2012), the authors summarized that the DMN, as one of the most widely studied resting-state networks, may be most significantly affected by chronic pain. In (Qi et al., 2016a), the study found that the average functional connectivity of the DMN in IBS patients was negatively correlated with IBS-SSS (IBS symptom severity scale), which is a scoring system used for assessing the severity of symptoms in patients with IBS. Compared to the healthy control group, the functional connectivity between DMN subregions in IBS patients was weakened, partly explaining the phenomenon of disordered visceral sensation in IBS patients. It is believed that chronic pain induced by IBS does indeed alter the DMN.

Among the top-ranking brain regions, the Inferior Parietal Lobule (IPL.R) is a part of the parietal lobe of the brain, primarily involved in processing sensory information and responding to external stimuli. This brain region is primarily engaged in attention and emotion regulation processes and is also associated with the regulation of the autonomic nervous system. This study (Skrobisz et al., 2019) suggests that gastrointestinal disturbances caused by Irritable Bowel Syndrome (IBS) may lead to lowered anxiety and pain thresholds, causing patients to focus more on negative outcomes. Therefore, we posit that changes in IPL.R, as a key brain region involved in attention and emotion regulation processes, due to the influence of IBS are understandable. In addition to the Inferior Parietal Lobule (IPL.R), the Inferior Frontal Orbital part (ORBinf.R), Middle Frontal Orbital part (ORBmid.R), and Superior Medial Frontal Orbital part (ORBsupmed.L) all belong to the frontal lobe of the brain and play important roles in emotion regulation, with the latter two being closely related to social behavior. In this study (Skrobisz et al., 2019), researchers found that psychological and social factors significantly influence the mechanisms regulating visceral sensitivity in the brains of IBS patients. The Postcentral Gyrus (PCG.R) is a part of the parietal lobe of the brain and is involved in the processing of bodily sensations. The study (Van Oudenhove et al., 2020) discovered that this brain region shows significant activation during visceral stimulation, indicating a higher likelihood of alterations under the chronic effects of IBS. The Postcentral Lobule (PCL.R) is responsible for sensory-motor control, body positioning, and certain higher cognitive functions. The Superior Frontal Orbital part (ORBsup.R) is located above the frontal lobe and also has a significant impact on emotion cognition and regulation. Through meta-analysis of previous related studies, a review (Yu et al., 2022) revealed that in IBS patients, the brain regions showing activation or deactivation are mainly associated with visceral sensation (PoCG, ACC, INS), emotional processing (hippocampus) and pain processing (frontal lobe, parietal lobe, supplementary motor area (SMA), thalamus, hippocampus, cerebellum, caudate nucleus, and PCUN). Our study also found that patients with IBS exhibit related anomalies in a wide range of frontal and parietal brain regions. From the overview of the aforementioned top 7 ranked brain regions, it is evident that these regions are mainly located in the parietal and frontal cortical areas of the brain, which is also confirmed by the review (Yu et al., 2022). One of the essential functions of the parietal cortical areas is the processing of sensory information, while the frontal lobe is a crucial region involved in emotion regulation. Therefore, we have reason to believe that the results obtained from the interpretability module are indeed strongly correlated with a wide range of frontal and parietal regions, which is well aligned with the prior IBS findings.

Next, we explore the brain regions that are ranked slightly lower in our interpretable output list. Through the summary of previous related research, we observed that these brain regions, such as the caudate nucleus, anterior and posterior cingulate gyrus, supplementary motor area, thalamus, and hippocampus, have been mentioned in previous studies multiple times, thereby validating the validity of the interpretability module. For example, in (Weng et al., 2017), the study used functional connectivity density (FCD) to investigate changes in the overall brain functional connectivity patterns in IBS patients. The experimental results showed that compared to the healthy control group, IBS patients showed reduced long- and short-range FCD in the bilateral anterior cingulate cortex (aMCC), decreased short-range FCD in the caudate nucleus, and increased long-range FCD in the right supplementary motor area. In (Qi et al., 2016b), using multiple covariate regression, it was found that patients with IBS showed higher positive resting-state functional connectivity between the amygdala and the adjacent hippocampus, as well as motor areas. In (Li et al., 2021a), the authors demonstrated that IBS patients exhibited abnormal functional connectivity in brain regions associated with the frontal-insular system and the somatosensory-motor network, especially the insular cortex and supplementary motor area (SMA), explaining the vicious cycle between negative emotions and gastrointestinal symptoms in IBS. In (Aziz et al., 2021), abnormal brain regions were observed in functional imaging of IBS patients, such as cortical thinning in the anterior cingulate cortex and increased gray matter in the thalamus. In (Ao et al., 2021), a voxel-based analysis of the amplitude of low-frequency fluctuation (fALFF) in the fALFF map of IBS and HC was compared, revealing changes in fALFF values in the left hippocampus. In (Ke et al., 2015), the analysis of regional homogeneity (ReHo) in patients showed an increase in ReHo in the thalamus and a decrease in ReHo in the anterior cingulate cortex compared to the control group. In (Mao et al., 2020), it was found that resting-state functional connectivity between the right habenula and the right thalamus was significantly reduced in IBS patients.

In addition to validation through prior literature review, we conducted an external validation to further confirm the efficacy of the interpretability module. For this, we utilized the top 20 regions identified by the whole brain model to predict IBS, comparing the outcomes against those obtained using 20 randomly selected brain regions. The findings demonstrated that the chosen 20 regions significantly outperformed the random regions in accurately classifying IBS and control participants, suggesting a more pronounced connection between the selected regions and IBS abnormalities.

Furthermore, we conducted PBM and correlation analyses on the brain regions identified by the interpretability module. For the correlation analysis results, we analyzed them sequentially based on the different attributes of the nodes. The degree of a node indicates the number of edges connected to that node, a higher degree suggests that the node has more connections with other nodes, potentially representing greater influence or higher levels of activity. According to the results of the correlation analysis, the IPL.R, which is involved in processing sensory information and responding to external stimuli, primarily participates in attention and emotional regulation processes. The subsequent regions, including ORBinf.R, ORBmid.R, and ORBsupmed.L, are also critical areas for emotional regulation. Notably, the above-mentioned brain regions exhibited a significant positive correlation with the IBS-SSS scores, indicating that increased activity in these regions is associated with greater severity of symptoms in IBS patients. The clustering coefficient of a node measures the tightness of connections among its neighbors, a high clustering coefficient suggests that the neighbors of a node are likely interconnected. Within this node attribute context, ORBinf.R, ORBmid.R, and PCG.R, which show a significant positive correlation with IBS-SSS, are important regions for emotional regulation in the brain. In contrast, IPL.R, CAU.R, and ACG.R are involved in visceral perception and pain perception, suggesting that the tighter the connections between these brain areas and surrounding regions, the higher the severity of symptoms in IBS patient.

### Limitations and future directions

While ST-GCN has greatly increased the predictive power of IBS classification using brain fMRI data, it is still insufficient to fully utilize the rich information contained within the data. Specifically, concerning the ST-fMRI-GCN model, we hold the view that its handling of temporal features is overly simplistic, missing out on the opportunity to fully leverage the dynamic correlations present in fMRI data. In future research, we believe that the following aspects can be improved: (1) Our study lacks longitudinal trials covering the progression of IBS. Longitudinal studies are necessary to track neural changes from different stages, allowing for a better understanding of how the onset and progression of IBS affect the organization of large-scale brain networks. (2) In the ST-fMRI-GCN, the model parameters, such as window size, spatial scale and temporal scale, are manually selected. Although numerous experiments have been conducted to maximize the classification accuracy, this method consumes a significant amount of time and manpower. In future experiments, we aim to improve the model by integrating methods that enable it to adaptively choose its hyperparameters, streamlining the optimization process. (3) In our interpretability module, we can identify brain regions that have been altered by IBS, but we cannot discover changes in brain functional connectivity influenced by IBS, nor can we determine if the altered brain regions are more active or inhibited. In future research, we aim to refine and enhance the interpretability module to improve its functionality. (4) Currently, despite the fact that the sample size of the IBS dataset utilized in our study significantly exceeds that of prior research, it appears challenging to meet the data requirements necessitated by deep learning models, and increase the risk of overfitting. This dataset has been collected over a span of more than three years and is still in the process of ongoing collection. We anticipate the opportunity to evaluate our model on a larger dataset in the future. (5) Due to the privacy concerns associated with IBS data, we were only able to obtain data from the collaborating hospital, which has limited the validation of the model’s generalizability and applicability. In future efforts, we intend to seek additional collaborative hospitals to establish a multi-site dataset that will facilitate the validation of our model.

## Conclusion

In this paper, we apply ST-GCN to our own IBS dataset, achieving the highest average classification accuracy of 83.51%. After the model training, we deduce through the interpretability module that brain regions such as the IPL.R, ORBinf.R, PCG.R, ORBmid.R and ORBsupmed.L are highly correlated with the classification of this disease.We found that multiple brain regions, such as ORBinf.R, PCG.R, ORBmid.R, were involved in the regulation of emotions. Most of these regions are located in the parietal and frontal lobes of the brain, with many of them belonging to the DMN.

## Acknowledgments

This work was supported in part by the Shenzhen Science and Technology Innovation Program under Grant JCYJ20220818102414031. This work was also supported by National Natural Science Foundation of China (No. 82071992)

## Conflict of Interest

None.

During the preparation of this work the author(s) used chatGPT in order to translation. After using this tool, the authors reviewed and edited the content as needed and take full responsibility for the content of the publication.

## Notes

### Competing Interest Statement

The authors have declared no competing interest.

